# The mutation rate of SARS-CoV-2 is highly variable between sites and is influenced by sequence context, genomic region, and RNA structure

**DOI:** 10.1101/2025.01.07.631013

**Authors:** Hugh K. Haddox, Georg Angehrn, Luca Sesta, Chris Jennings-Shaffer, Seth D. Temple, Jared G. Galloway, William S. DeWitt, Jesse D. Bloom, Frederick A. Matsen, Richard A. Neher

## Abstract

RNA viruses like SARS-CoV-2 have a high mutation rate, which contributes to their rapid evolution. The rate of mutations depends on the mutation type (e.g., `A`→`C`, `A`→`G`, etc.) and can vary between sites in the viral genome. Understanding this variation can shed light on the mutational processes at play, and is crucial for quantitative modeling of viral evolution. Using the millions of available SARS-CoV-2 full-genome sequences, we estimate rates of synonymous mutations for all 12 possible nucleotide mutation types and examine how much these rates vary between sites. We find a surprisingly high level of variability and several striking patterns: the rates of four mutation types suddenly increase at one of two gene boundaries; the rates of most mutation types strongly depend on a site’s local sequence context, with up to 56-fold differences between contexts; consistent with a previous study, the rates of some mutation types are lower at sites engaged in RNA secondary structure. A simple log-linear model of these features explains ∼15-60% of the fold-variation of mutation rates between sites, depending on mutation type; more complex models only modestly improve predictive power out of sample. We estimate the fitness effect of each mutation based on the number of times it actually occurs versus the number of times it is expected to occur based on the model. We identify several small regions of the genome where synonymous or noncoding mutations occur much less often than expected, indicative of strong purifying selection on the RNA sequence that is independent of protein sequence. Overall, this work expands our basic understanding of SARS-CoV-2’s evolution by characterizing the virus’s mutation process at the level of individual sites and uncovering several striking mutational patterns that arise from unknown mechanisms.

## Introduction

SARS-CoV-2 evolves rapidly. In the four years since the virus emerged, dozens of mutations have risen to high frequencies in the global viral population, including ones that increase its transmissibility and help it evade human immune responses [1, 2, 3, 4, 5, 6]. As it transitions to becoming endemic in humans, it continues to evolve resistance to updated immune responses [7].

One factor that shapes SARS-CoV-2’s evolution is its mutation rate (i.e., the rate of mutations in the absence of selection). Its mutation rate is not as high as many other RNA viruses, due to a proofreading mechanism present in coronaviruses and other members of the *Nidovirales* order [8, 9]. However, SARS-CoV-2’s mutation rate is still orders of magnitude higher than that of cellular organisms [10], providing fuel for the virus’s rapid evolution. SARS-CoV-2’s mutation rate is also variable. Studies quantifying the rates of each of the 12 possible nucleotide mutation types (`A`→`C`, `A`→`G`, etc.) found that the rates span two orders of magnitude [11, 12]. The rates of a few mutation types were also found to have shifted over evolutionary time, adding additional variability [12, 13].

This high variability in rates can bias which paths SARS-CoV-2’s evolution is mostly likely to take. For instance, the nucleotide mutation type `C`→`T` has the highest neutral mutation rate [11, 12]. As might be expected from this, amino-acid mutations arising from `C`→`T` mutations are much more common than amino-acid mutations arising from other mutation types [14, 15]. Thus, it is important to identify factors that modulate the virus’s mutation rate.

An open question is how much the rates of the 12 nucleotide mutation types vary between sites in the SARS-CoV-2 genome. RNA-virus mutations arise from several mechanisms, including misincorporations by the viral polymerase, components of the innate anti-viral immune response of the host, and reactive oxygen species. Some mechanisms are highly specific to a particular mutation type. The very high `C`→`T` mutation rate of SARS-CoV-2 has been hypothesized to be due to deamination through host enzymes [14, 16, 15], while the high `G`→`T` rate has been hypothesized to be due to oxidative damage [13]. These editing mechanisms are known to depend on the target site’s local sequence context and physical accessibility, and, unlike misincorporations, their rates are expected to be strand-specific. Thus, we expect the resulting mutation rates to vary between sites.

Past studies have found evidence that SARS-CoV-2’s mutation rate varies between sites, with one study finding that mutation rates depended weakly on a site’s local sequence context (though this study only examined a few mutation types) [11], and others finding that the rates of some, but not all, mutation types strongly depended on RNA structure [17, 18]. Other studies have modeled differences in mutation rates between sites [19, 20], but did not separately examine individual mutation types. Thus, we still lack a comprehensive picture of how the neutral rates of the 12 mutation types vary between sites, and to what extent sequence features such as local context and RNA structure explain this variability.

Through an impressive effort by many researchers around the globe, we now have ∼16 million full-length SARS-CoV-2 viral genome sequences (each a consensus sequence from an infected human host)—orders of magnitude more than for any other virus [21]. Further, these sequences were all collected in a ∼ 4-year period, providing very dense sampling of the evolutionary history of SARS-CoV-2; in a phylogenetic tree using all these sequences, branches typically have only one or two mutations, corresponding to just a few weeks of viral evolution [22, 23]. This implies that observed mutations with small fitness effects are only weakly affected by natural selection and can be used to estimate the neutral mutation rate [24].

Some studies have used the above data to estimate the rates of the 12 nucleotide mutation types as genome-wide averages [11, 12, 25]. However, a recent study by Bloom and Neher showed that the SARS-CoV-2 data are rich enough to also estimate evolutionary rates at the level of individual sites in the genome [24]. By comparing rates of mutations that change amino acids in viral proteins to aggregate rates of synonymous mutations that provide a neutral baseline, the study also estimated fitness effects of amino-acid mutations at nearly all coding sites in the genome, providing a highly detailed picture of the selective pressures acting on viral proteins. However, the study did not examine how much estimated neutral rates of the 12 nucleotide mutation types vary between sites, nor attempt to model these rates in terms of sequence characteristics.

In this paper, we show that these rates are highly variable between sites, with the 5th and 95th percentiles of rates for a given mutation type differing up to ∼100-fold. We find that a site’s rate depends on features related to its genomic region, RNA structure, and local sequence context. The manner in which each feature does so is highly idiosyncratic between mutation types. Depending on the mutation type, our model that includes these features explains between ∼15-60% of the fold-variation of mutation rates between sites. This suggests that we have identified some, but not all, of the features that contribute to this variation. We use this model to explore signatures of selection on synonymous and noncoding mutations along the genome. We also use the model to update fitness-effect estimates for amino-acid mutations reported by Bloom and Neher [24], which previously did not account for the effect of sequence context on a site’s expected neutral mutation rate. We make the updated fitness estimates openly available at https://github.com/neherlab/SARS2-mut-fitness-v2.

## Results

### Estimating site-specific synonymous mutation rates for each mutation type

We sought to estimate SARS-CoV-2’s mutation rate at the level of individual sites in the genome, doing so for each of the 12 possible nucleotide mutation types. We used a modified version of the approach from De Maio et al. [11] and Bloom et al. [12]. For illustrative purposes, we describe how this approach works for a single mutation type, `T`→`G` (Figure 1A). Using the Wuhan-Hu-1 genome as a reference sequence, we first identified all nucleotide sites that meet the following criteria: the reference nucleotide is a `T`; a `T`→`G` mutation is synonymous in all open reading frames that contain the site; and the site, as well as the nucleotides that immediately border it or that are in the same codon, are highly conserved (see *Methods*). Next, for each of these sites, we counted the number of occurrences of a synonymous `T`→`G` mutation at that site along the branches of an UShER phylogenetic tree with ∼16 million SARS-CoV-2 genome sequences from the GISAID database [21, 22, 23]. Note that this approach is not the same as counting the abundance of mutations among the ∼16 million leaves of the tree: if a mutation occurs on a branch that gives rise to many descendants carrying that mutation, the mutation event on the branch is counted just once. We obtained mutation counts from the computational pipeline of Bloom and Neher [24], which imposes several quality-control filters that help eliminate errors in the data. The resulting mutation counts per site provide an estimate of relative persite `T`→`G` substitution rates (Figure 1B). If the effects of synonymous mutations are assumed to be neutral, then these *substitution* rates correspond to neutral *mutation* rates. Below, we investigate where the assumption that synonymous mutations are neutral might break down.

**Figure 1:**
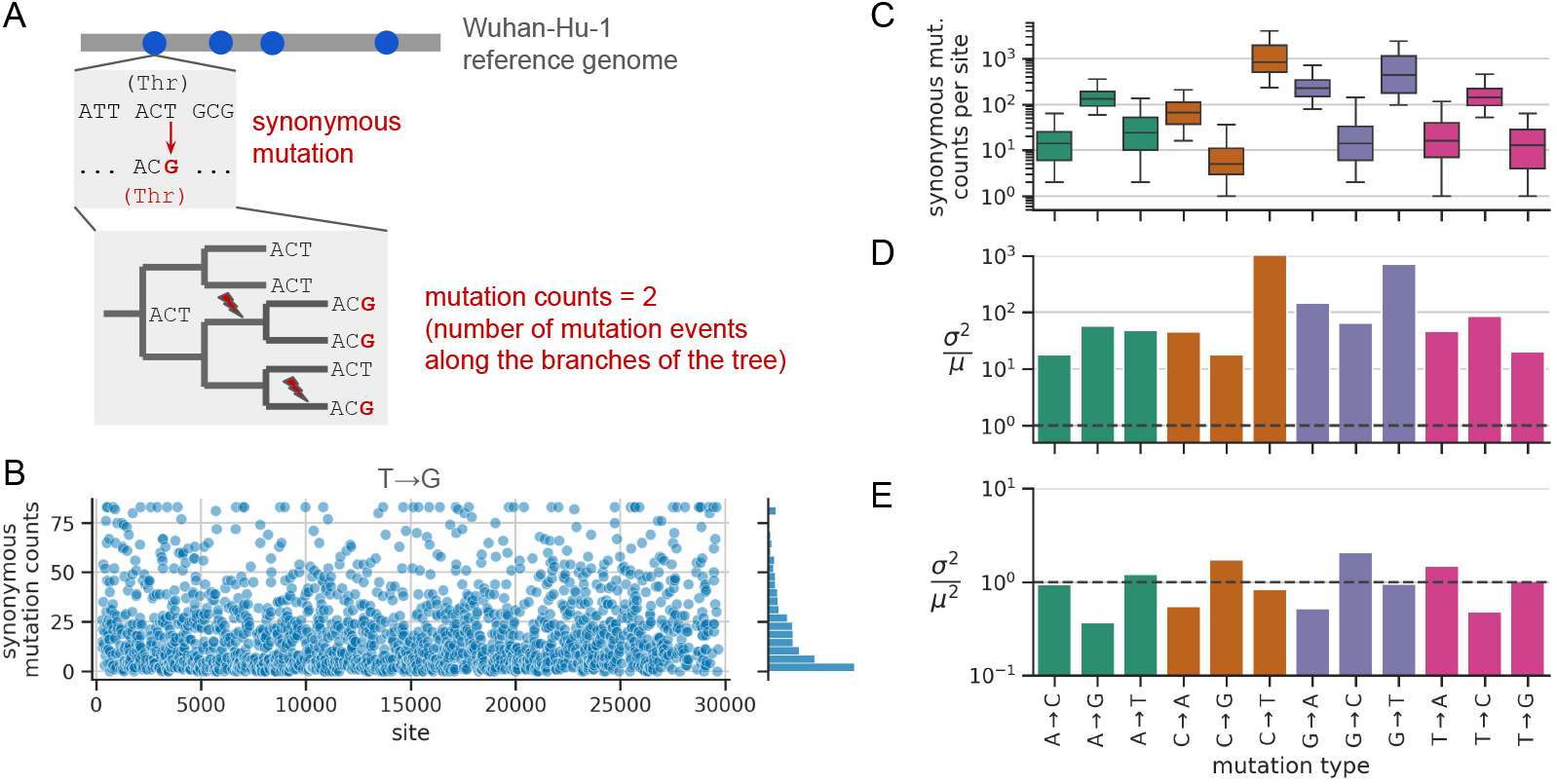
Estimation of site-specific mutation rates. **(A)** Estimating site-specific mutation rates, illustrated for `T`→`G` mutations. First, we identified all sites (blue dots) where the reference nucleotide is a `T`, a `T`→`G` mutation is synonymous, and the site and its local sequence context are highly conserved. Next, we counted the number of synonymous `T`→`G` mutation events on the branches of an UShER tree. The resulting site-specific counts are proportional to rates. We separately implemented this strategy for each of the 12 possible mutation types. Figure adapted from Bloom and Neher [24]. **(B)** Site-specific synonymous mutation counts for `T`→`G` mutations computed using the workflow from panel A, with a ceiling applied to the distribution at its 98th percentile, and plotted as a function of nucleotide site in the SARS-CoV-2 genome. For each mutation type, **(C)** shows the distribution of site-specific synonymous mutation counts across those sites (whiskers show the 5th and 95th percentiles); **(D)** shows the variance of each distribution divided by the distribution’s mean (after applying a ceiling to the distribution at its 98th percentile), with the dashed line at 1.0 showing the value expected for Poisson-distributed data; and **(E)** shows the variance of each distribution divided by the distribution’s mean squared (these values are more uniform and cluster near 1.0).

By only analyzing mutations at sites that are highly conserved, we restrict the analysis to mutations that are possible on nearly all branches of the tree. This strategy helps ensure that differences in mutation counts between sites are proportional to differences in mutation rates. In contrast, at sites that are *not* conserved, differences in mutation counts between sites can also strongly depend on evolutionary opportunity (e.g., some mutations are only possible on a small fraction of branches of the tree, which would tend to dramatically reduce the counts of these mutations relative to the mutations in our analysis). By only analyzing sites with local sequence contexts that are highly conserved, we isolate the effect of local sequence context on mutation rate. All genome sequences in our dataset are >99% identical to the Wuhan-Hu-1 reference sequence, and so most sites are highly conserved. We applied the above approach to each mutation type, obtaining counts data for 100s to 1,000s of sites per mutation type (Table S1). As expected from previous analyses of the mutational spectrum and nucleotide composition of the SARS-CoV-2 genome, mutations away from `T` (we indicate mutations away from `T` to any nucleotide by `T` →`N`), tend to have the most sites with data, followed by mutation types `A`→`N`, `C`→`N`, and `G`→`N` [12]. Figure 1C shows the distribution of synonymous mutation counts per site for each mutation type. The median counts per site vary from less than 10 for `C`→`G` mutations to nearly 1,000 for `C`→`T` mutations. While previous studies have primarily used this type of counts data to estimate differences in genome-wide mutation rates between mutation types, we sought to analyze variability in rates between individual sites.

### Rates of synonymous mutations are highly variable between sites

While we expected some level of variability in synonymous mutation counts between sites for a given mutation type, the variability that we observed was surprisingly high. For `T`→`G` mutations, although many sites had zero or very low counts, an appreciable number of sites had counts of 50 or higher (Figure 1B). The other mutation types also had broad distributions, with the 5th and 95th percentiles of most distributions spanning one to two orders of magnitude (Figure 1C).

The observed variability is much higher than expected from statistical sampling noise. If all sites had the same underlying mutation rate for a given mutation type and differences in counts between sites were purely due to statistical noise, then it is reasonable to expect the counts are Poisson-distributed around a common mean that is equal to the distribution’s variance. However, the counts are substantially overdispersed: the variance is ∼10-1,000 times larger than the mean, depending on mutation type (Figure 1D). We find that the variance is roughly equal to the mean squared (Figure 1E), with the counts distributions closely resembling a log-normal distribution (Fig. S1). This suggests that the counts arise from factors with multiplicative effects. Below, we explore the explanatory power of several sequence features in a multiplicative model.

Differences in counts between sites might arise from experimental errors and computational artifacts used to generate the sequencing data. Errors have been found to dramatically inflate counts at a small subset of sites [26, 27]. As such, we omit sites known to be error-prone. As a control, we computed site-specific counts of premature stop mutations in essential genes, which are highly deleterious. Nearly all such mutations have very low counts (Fig. S2). Since sequencing errors are expected to occur regardless of biological effects of a mutation, the very low counts of premature stop mutations suggests that the counts are not dominated by errors.

In the next three sections, we identify factors that are associated with differences in mutation rates between sites. To inform this effort, we first sought to determine if site-specific rates were correlated between mutation types with the same wildtype nucleotide (e.g., among sites where the wildtype nucleotide is `T`, and both `T`→`G` and `T`→`C` mutations are synonymous). For most pairs of mutation types, the correlation is low (*R*^2^ *<* 0.1; Fig. S3), suggesting that variability in site-specific counts often depends on factors that are specific to a particular mutation type. This result motivated us to analyze each mutation type separately.

### Rates of synonymous mutations depend on genomic region

First, we examined whether synonymous mutation counts vary systematically along the genome on large scales. To do so, for each mutation type, we computed the median synonymous mutation counts in a 2-kilobase sliding window across the genome. In this and other analyses, we use the median instead of the mean since the mean can be skewed by a small number of outlier sites with very high counts. Figure 2A-C show the sliding-window median for three example mutation types, while Figure 2D summarizes the traces for all 12 mutation types. For some mutation types, like `A`→`G`, the sliding-window median was roughly uniform across the genome (Figure 2A). In contrast, a few other mutation types showed strikingly non-uniform patterns.

**Figure 2:**
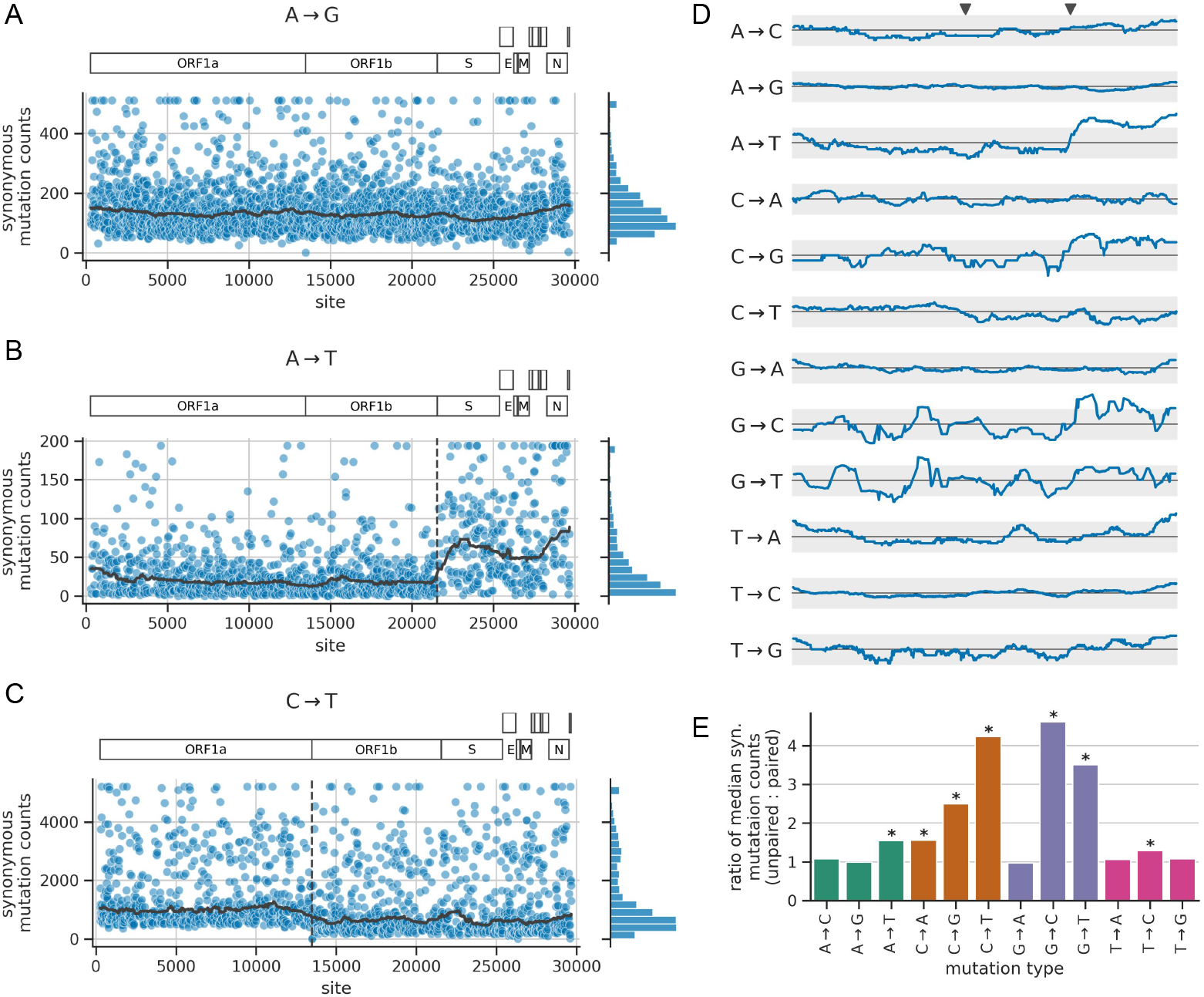
Distribution of mutation counts as a function of genomic region and RNA secondary structure. **(A)-(C)** Each scatter plot shows site-specific synonymous mutation counts for a given mutation type, applying a ceiling at the counts distribution’s 98th percentile. The bold gray line shows the median counts in a 2-kilobase sliding window across the genome. The vertical dashed lines show gene boundaries that correspond with abrupt changes in some counts distributions. **(D)** Traces of sliding-window median synonymous mutation counts for each mutation type, similar to those from panels A-C. The x-axis spans the length of the genome, with the two gray triangles above the top plot showing the location of the gene boundaries highlighted in panels B and C. The y-axis shows the ratio of each sliding-window median relative to the genome-wide median, with the solid horizontal black line showing a ratio of 1.0 and the gray region spanning ratios of 0.5 and 2.0. Ratios are plotted on a log scale, such that the gray region deviates from the solid line by the same amount in either direction. **(E)** Ratio of median synonymous mutation counts at sites that are unpaired to sites that are base-paired in the predicted secondary structure of the SARS-CoV-2 genome. An asterisk indicates that the ratio for a given mutation type is significantly >1.0, as determined by randomization testing (see *Methods*), using a threshold of p*<*0.05 after Bonferroni correction for multiple-hypothesis testing.

The mutation type `A`→`T` showed the strongest and sharpest dependence of mutation rate on genomic region (Figure 2B). At the boundary between the ORF1b and spike genes, the median counts suddenly increases by more than a factor of two. The mutation types `C`→`G` and `G`→`C` show a similar, but weaker pattern (Figure 2D; Fig. S4). Of note, `T`→`A` mutations do *not* show the pattern, which suggests the mechanism giving rise to the `A`→`T` pattern is strand dependent. Indeed, the mutation counts are all computed relative to the nucleotide sequence of the virus’s positive-sense genomic RNA; if the `A`→`T` mutation process operated the same way on both strands, then we would also expect to see the above pattern for the complement mutation type, `T`→`A`.

Since the genomic region with elevated mutation counts overlaps with regions encoded by subgenomic RNAs [28], we wondered whether many of the mutations uniquely derive from subgenomic RNAs, rather than full-length genomic RNAs. There are two pieces of evidence against this hypothesis. First, the mutation counts are derived from consensus sequences, and rare mutations in subgenomic RNAs would not change the consensus sequence. Second, the `A`→`T` pattern is present both for mutations on terminal branches and mutations deeper in the tree (Fig. S5). Subgenomic RNAs are not vertically transmitted between viruses, so mutations to those RNAs would be expected to be concentrated on terminal branches rather than deeper branches.

A few other mutation types show notable, but less pronounced patterns. For `C`→`T` mutations, the floor of synonymous mutation counts per site suddenly decreases near the end of ORF1a close to the ribosomal slippage site separating ORF1a from the rest of the genome (Figure 2C). And for some mutation types, the counts gradually increase as the sliding window approaches either end of the genome (Figure 2D; the pattern is most pronounced for `T`→`A`). The sliding-window median is noisier for mutation types that have fewer sites with data (e.g., `C`→`G`, `G`→`C`, and `G`→`T`), and so the differences in median are not well-supported by the data. In all, synonymous mutation counts depend on genomic region for some, but not all mutation types, and do so in an idiosyncratic manner.

### Rates of synonymous mutations depend on RNA secondary structure

Previous studies found that the rates of certain mutation types are lower in regions of the SARS-CoV-2 genome that are predicted to be involved in RNA secondary structure [17, 18]. The study by Hensel used a dataset of synony-mous mutation counts that is similar to the one we generated here [18]. Hensel considered a secondary-structure model of the SARS-CoV-2 genome predicted from dimethyl-sulfate reactivity data that chemically probes for structure at the level of individual sites [29]. In this structure, roughly half of all sites are predicted to be base-paired with another site, while the other half are predicted to be unpaired. Hensel found that synonymous mutation counts were significantly higher at unpaired sites compared to paired sites for 7 of the 12 mutation types. However, Hensel only quantified effect sizes (i.e., how much counts differ between unpaired and paired sites) for 4 of the 7 significant mutation types. We sought to reproduce the above findings and to quantify effect sizes for all significant mutation types.

In our data, the ratio of median synonymous counts between unpaired vs. paired sites ranges from about 1 to 4 depending on mutation type (Figure 2E). This ratio is significantly >1 for the same 7 mutation types identified by Hensel. These ratios quantify effect sizes for each mutation type, including the 3 significant mutation types not quantified by Hensel: `A`→`T`, `C`→`G`, and `G`→`C`, the latter of which has the highest ratio of any mutation type. The effects are not mirrored by complement mutation types, providing another example of strand asymmetry. `C`→`T` mutations were unique in that they displayed a clear bi-modal rate distribution where the higher-rate peak consists of sites predicted to be unpaired and the lower-rate peak coming from sites predicted to be paired, suggesting that pairing is an especially strong determinant of `C`→`T` mutation rate (Fig. S6). Overall, the rate of some but not all mutation types depends strongly on RNA structure in a strand-specific manner.

### Rates of synonymous mutations depend on local sequence context

To investigate if site-specific synonymous mutation counts depend on local sequence context, we grouped sites by the 3-mer nucleotide motif centered on a given site and quantified differences in counts between motifs. We performed this analysis separately for each of the 12 mutation types. To limit the impact of noise, we only analyzed motifs with positive counts at 10 or more sites.

Most mutation types show large differences in counts between 3-mer motifs. The `T`→`G` mutation type shows the most striking differences, where the median synonymous mutation count per motif range from 1 to 56 (Figure 3A). The process behind the motif dependence of `T`→`G` mutations appears to be symmetric between strands, as the median `T`→`G` counts in a given motif closely mirrors the median `A`→`C` counts in the corresponding reverse-complement motif (Figure 3B).

**Figure 3:**
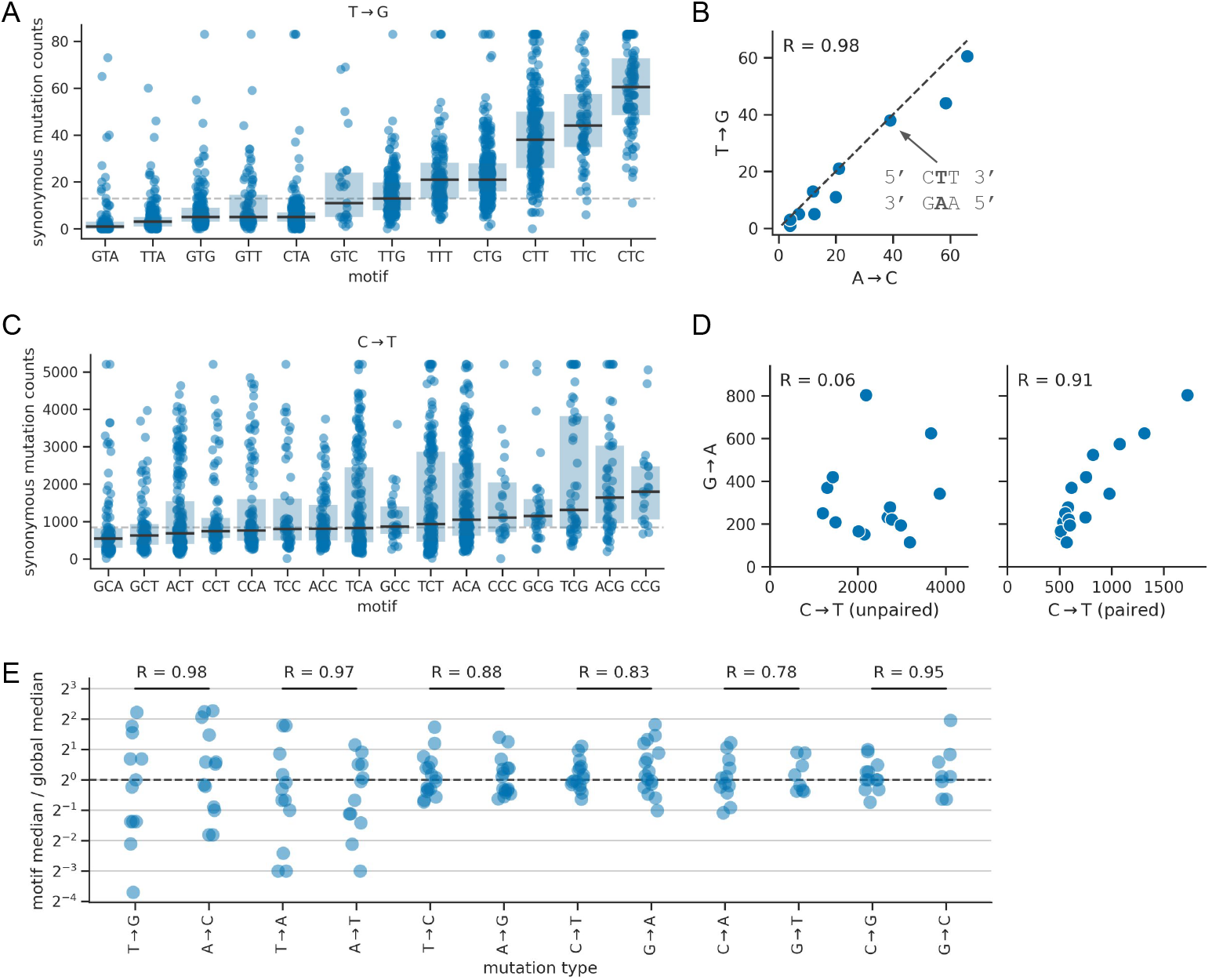
Synonymous mutation counts depend on local sequence context, but the strength of the effect depends on mutation type. **(A)** Counts for `T`→`G` mutations, where each dot corresponds to a site where a `T`→`G` mutation is synonymous, and where sites are grouped on the x-axis by the 3-mer sequence motif centered on the site. Box plots show the median and interquartile range of each distribution. The horizontal dashed line shows the global median counts of all sites. We applied a ceiling to the counts distribution at the 98th percentile of all sites. **(B)** Correlation of median counts of synonymous `T`→`G` mutations in a given 3-mer motif with median counts of synonymous `A`→`C` mutations in the reverse complement of that motif. The text attached to the arrow highlights an example reverse-complement motif pair. The data are highly correlated (R is the Pearson correlation coefficient) and track the line of equivalence (diagonal dashed line). **(C)** Same as panel A, but with data for `C`→`T` mutations, which is an example of a mutation type where local sequence context has a more modest effect on counts. **(D)** Same as panel B, but with data for `C`→`T` and `G`→`A` mutations and performing the analysis separately for `C`→`T` data based on the RNA secondary structure at a site (paired or unpaired). These plots do not show the line of equivalence as the counts do not show a 1:1 ratio. **(E)** Summary of patterns across all mutation types. For a given mutation type, each dot corresponds to a 3-mer motif. Its value on the y-axis is the fold difference between the median synonymous mutation counts of sites in that motif and the global median across all sites from all motifs. On the x-axis, each mutation type is plotted next to its complement mutation type, with horizontal black bars connecting pairs, and the value above each bar reporting the Pearson correlation coefficient (R) of median synonymous counts between reverse-complement motif pairs, as shown in panels B (it does not divide sites into groups as in panel D).

In contrast, `C`→ `T` mutation rates are only modestly impacted by local sequence context (Figure 3C). Since `C`→`T` counts strongly depend on RNA secondary structure, we also analyzed paired and unpaired sites separately. We found that counts for both groups of sites depended on local sequence context, but with somewhat different motif dependencies (Fig. S7A). There is an interaction between pairing and whether the median `C`→`T` counts in a given motif correlated with the median `G`→`A` counts in the corresponding reverse-complement motif. Specifically, we found that the correlation was low at *unpaired* `C`→`T` sites and high at *paired* `C`→`T` sites (Figure 3D; `G`→`A` mutation counts did not depend on pairing). This could mean that the `C`→`T` mutational signature visible at paired sites comes from a process that operates on both strands—though not equally, as counts for `C`→`T` mutations are substantially higher than for `G`→`A` mutations—and that the `C`→`T` mutational signature visible at unpaired sites is largely strand specific. Thus, while the `T`→`G` mutational signature shows clear signs of strand symmetry, that of `C`→`T` shows more complicated patterns.

In general, for mutation types `A`→`N` and `T`→ `N`, we saw a strong dependence of counts on local motifs and symmetry between strands, with the transversion types (`A`→`C`, `A`→`T`, `T`→`A`, `T`→`G`) showing stronger dependence than the transition types (`A`→`G`, `T`→`C`; Figure 3E). In contrast, counts of `C`→`N` and `G`→`N` mutations were less dependent on local motifs. And while counts for these types are correlated among reverse-complement motifs, they do not show a 1:1 ratio. Instead, the counts are always higher for one mutation type in the complement pair, e.g. Figure 3D), indicating some degree of strand asymmetry. Not all mutation types had enough sites with data to confidently assess whether 3-mer motif patterns depend on genomic region or RNA secondary structure, as seen with `C`→`T` above. Of those with enough data, only `C`→`G`, `G`→`T`, and `C`→`T` showed substantial dependencies (Fig. S7).

Overall, these data suggest that local sequence context has a pronounced effect on a site’s mutation rate, with evidence for both symmetric and asymmetric mutation processes.

### Modeling context dependence of synonymous mutation rates

The above data exploration suggests that the synonymous mutation rate at site *i* depends on (a) the mutation type, (b) the site’s genomic region, (c) whether the site is base-paired in the genomic RNA secondary structure, and (d) the site’s local 5′ and 3′ nucleotide context. Anticipating that these factors modulate the mutation rate in a multiplicative manner, e.g. by modulation of an interaction energy with a mutagen or the accessibility of the base, we linearly regress (for all 12 mutation types separately) the natural logarithm of the observed mutation counts 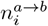 of type `a`→`b` at site *i* (including a pseudocount of *α* = 0.5). The indicator variable 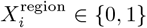, corresponding to factor (b), indicates site *i* is downstream from a given genomic boundary: the end of ORF1a for `C`→`T`; and the end of ORF1b for `A`→`T`, `C`→`G`, and `G`→`C` (we do not model this feature for other mutation types). The indicator variable 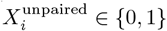, corresponding to factor (c), indicates *i* is predicted to be unpaired according to Lan et al. [29]. The indicator variables 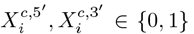 correspond to factor (d), so that 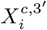 indicates the nucleotide at the 3′ context position *i* + 1 is *c* (analogously for the local 5′ context). These are indexed by the nucleotides *c* ∈{`C,G,T`}, so that the intercept term *β*^base^ is referenced to nucleotide `A` in both the 5′ and 3′ contexts. Our predictor takes the form:

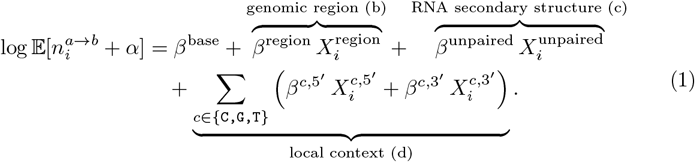

Note that all *β* coefficients are specific to the mutation type `a`→`b`, which we omit for brevity. We fit the model parameters using ridge regression by minimizing the sum of two terms: a loss term that quantifies the squared deviation between the observed and predicted log-counts and an L2 term that weakly penalizes non-zero *β* coefficients (excluding *β*^base^). The exponentiated fitted parameters *e*^*β*^ are the fold-changes in the mutation rate caused by turning on the corresponding *X*.

Depending on mutation type, the model that includes all three factors explains between 15-60% of the variance in the log-counts (Figure 4). Genomic region explains a significant part of the variance for each of the four mutation types it was used to model. RNA secondary-structure pairing state explains almost 35% of the variance of `C`→`T` mutations, while the local context explains a significant part of the variance for mutations away from `A` or `T` as well as for `G`→`A`. The decrease in explained variance from omitting a feature from the model closely mirrors how much variance the feature explains in the absence of other features (Fig. S8). This suggests that the contributions of each feature are largely orthogonal (i.e., the differences in counts explained by one feature cannot be explained by another). Furthermore, the explained variance does not decrease substantially when training the model on 80% of the data and validating it on the remaining 20% (see Fig. S9).

**Figure 4:**
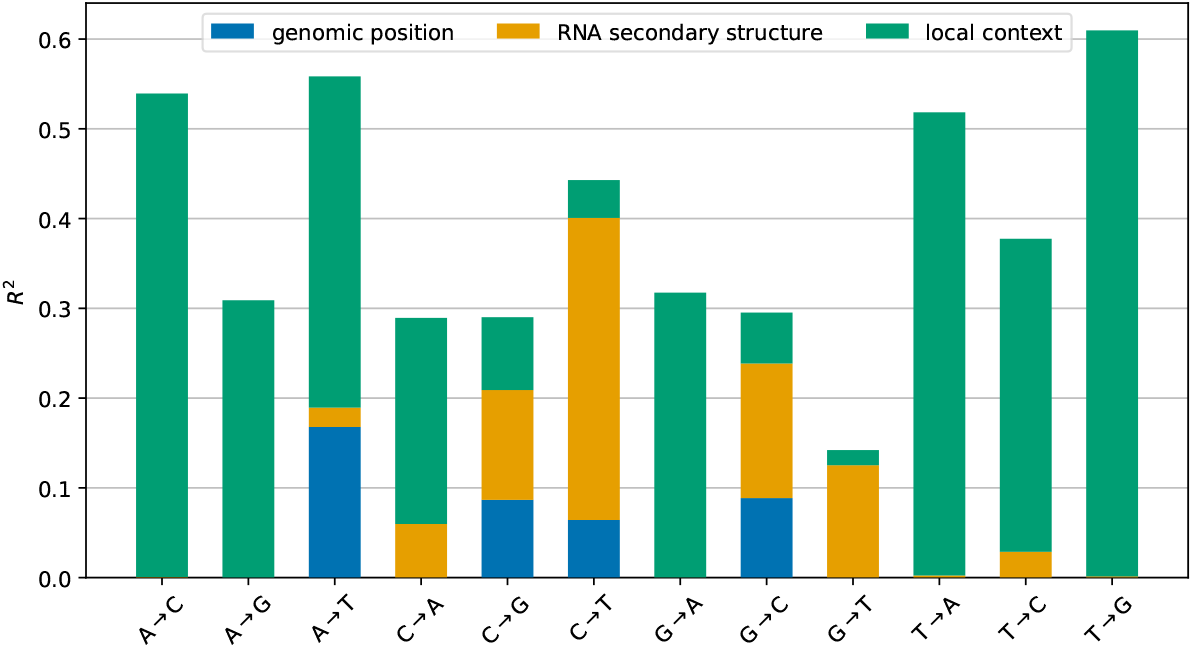
Explanatory power of different sequence features. For every mutation type, the graph shows the fraction *R*^2^ of the variance of the log-counts explained by the model as the features *genomic region, RNA secondary structure*, and *local context* are successively included in the log-linear model in Eq. (1). The model is trained and evaluated on the full dataset.

A limitation of the above strategy for modeling local sequence context is that it only considers the 5′ and 3′ nucleotides neighboring a site, and assumes additive effects of these sites on the logarithm of the mutation rate. This motivated us to explore more complicated models that consider wider sequence windows, and which allow for non-additive effects. An inherent challenge in doing so is the limited size of the training data relative to the number of parameters: while each mutation type only has 100s to 1,000s of sites with data, there are 256 possible 5-mer motifs per mutation type, making it difficult to separately model each motif. To address this challenge, we devised a predictive model that integrates information across sequence windows of a defined size using embeddings, but has a parameter-sparse architecture that reduces its propensity for overfitting (see *Methods*). We then separately fit a series of such models that consider windows of length 3, 5, and 7. For most mutation types, the models considering window lengths of 5 and 7 outperform those considering length 3 windows, as well as the above log-linear model, as quantified by their ability to predict counts at sites withheld from model training, with improvements in *R*^2^ of ∼0.1 (Fig. S9). This result suggests that the underlying mutational mechanisms depend on more than just the 5′ and 3′ nucleotides neighboring a site. However, for the three mutation types with the least training data, the more complex models tend to perform worse. Because of this, and because the log-linear model still captures most of the signal learned by the complex models, we elected to continue to use the easily interpretable log-linear model for the rest of our analysis.

### Purifying selection on synonymous and non-coding mutations is restricted to a few regions

As in Bloom and Neher [24], we have treated synonymous mutations as neutral and used them to estimate the neutral mutation rate. Any variation in their counts between sites was treated as a modulation of the underlying mutation rate rather than differences in functional constraints between sites. For this reasoning to be valid, only a minority of synonymous mutations should be under purifying selection, for example due to RNA secondary structure or regulatory elements (see below). We examined the distribution of fitness estimates along the genome to test this reasoning.

Following Bloom and Neher, we estimated the fitness effect of a mutation `a`→`b` at site *i* as the natural logarithm of the ratio of the number of observed counts of that mutation to the number of counts expected under neutral evolution:

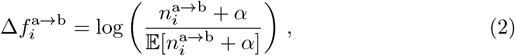

where *α* = 0.5 was used as a pseudocount to avoid invalid values. Bloom and Neher used the average counts of a given mutation type at 4-fold degenerate sites as the expected count, while we used the expected counts from the log-linear model described above. When Δ*f*_*i*_ is zero, this indicates that the mutation is neutral; positive or negative values indicate that the mutation count at this site is higher or lower than expected, which is interpreted as a beneficial or deleterious effect, respectively.

Fitness effects estimated using Eq. (2) directly depend on the accuracy of the estimated expected counts. We expect that estimates based on our log-linear model that captures context dependence will be more accurate than those by Bloom and Neher based on the average mutation counts at 4-fold degenerate sites. To assess accuracy, we examined the distribution of estimated fitness effects of synonymous mutations, most of which are expected to have neutral effects. This distribution is more tightly centered around zero when estimating effects using our revised expected counts (Figure 5A; the lower variance is a necessary consequence of the fact that we fit a model to reduce exactly this variation). The log-linear model is also less sensitive to outlier sites with extremely high mutation counts, which introduced a negative skew to fitness estimates by Bloom and Neher.

**Figure 5:**
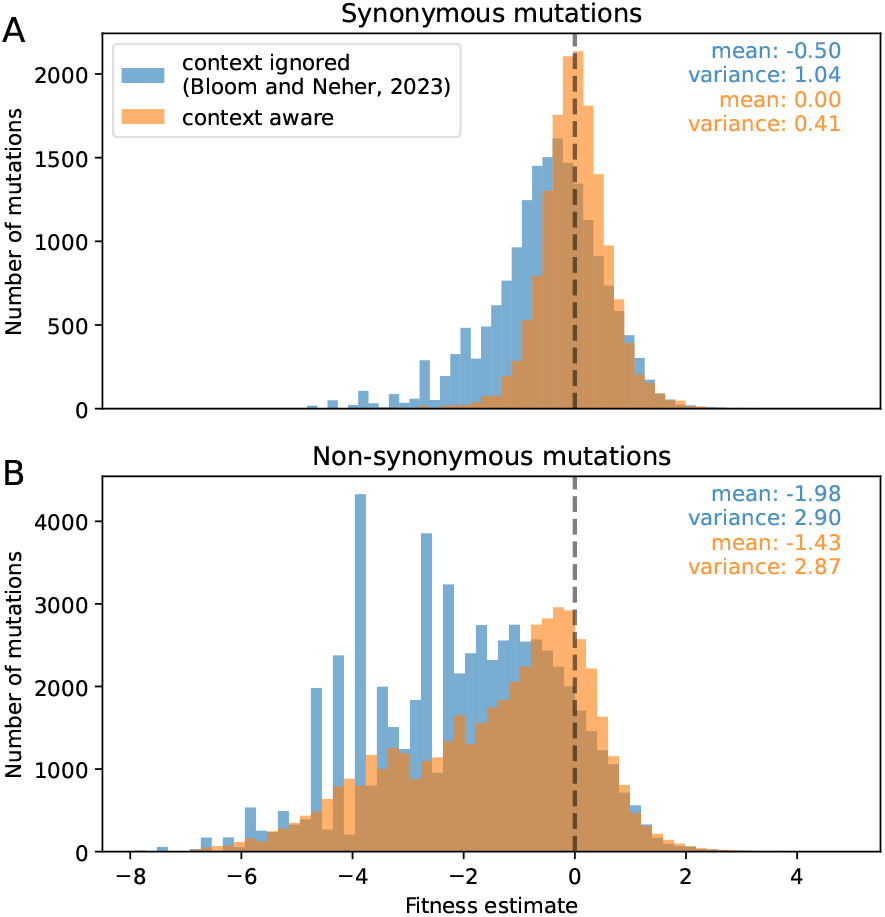
Improved fitness estimates. Accounting for the sequence context of mutations results in fitness estimates of synonymous mutations that are more tightly centered around 0 and removes artifactual peaks from the distribution of fitness effects of non-synonymous mutations. The histogram is restricted to mutations for which the predicted or expected counts are larger than 10.

When examining the distribution of fitness estimates of non-synonymous mutations, the expected counts by Bloom and Neher led to artificial peaks that are not present when using the expected counts from the log-linear model. These peaks were due to very deleterious mutations with observed counts equal to 0, which means the fitness estimate is determined by the pseudocount and expected count.

With the log-linear model of the expected counts, the variance of the distribution of non-synonymous fitness effects is 2.9, which is 7-fold larger than that of the synonymous effects (see Figure 5). Previously the difference was less than 3-fold, suggesting that the new model increases the contrast between the putatively neutral synonymous mutations and the non-synonymous mutations with a broad range of fitness effects.

Direct functional constraints of the RNA sequence, such as secondary structure elements or binding sites, manifests itself in purifying selection against synonymous mutations and in most cases we expect such constraint to affect several neighboring synonymous sites in the genome sequence. To identify such groups, we estimated a landscape of fitness effects of synonymous mutations across the genome, with the effects smoothed along neighboring sites (see Methods A.5). The smoothed fitness landscape will deviate strongly from zero only when fitness effects of mutations at several nearby sites are large. As expected, the smoothed effects hover around zero (Figure 6A), except for a few isolated areas where they dip substantially negative, which supports the idea that only a minority of synonymous positions are under strong purifying selection from such functional constraints. We repeated the analysis after permuting positions along the genome and did not see such pronounced peaks (gray lines in Figure 6A&B).

**Figure 6:**
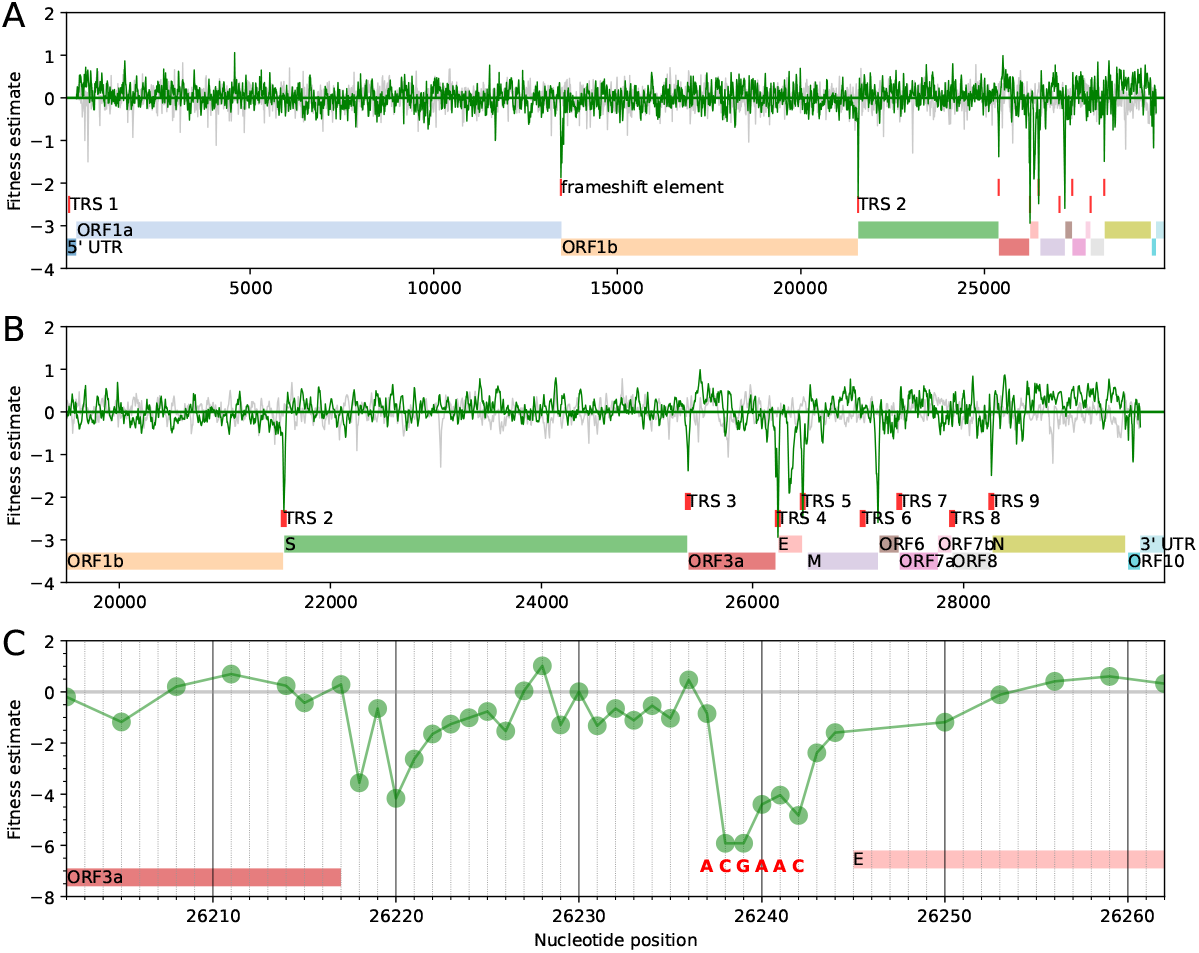
Synonymous mutations are under purifying selection at a few regions in the genome. Fitness estimates for synonymous mutations as defined in Eq. (2) along the full SARS-CoV-2 genome (A), along the structural and accessory genes (B), and around transcription regulatory site 4 (TRS 4) (C). The transcription regulatory sites were identified by marking the TRS motif ACGAAC in the reference genome. Note that panels (A) and (B) show a version of the fitness estimate that was smoothed along the genome as detailed in the Methods section. The faint gray line in A&B shows the fitness estimates after permuting the positions along the genome. For panel (C), the mutation counts of all non-excluded synonymous or non-coding mutations at a site were aggregated to obtain one estimate per site. ORF = Open Reading Frame, S = Spike, E = Envelope, M = Membrane, N = Nucleoprotein.

Most isolated regions where the fitness effects are substantially negative correspond to well-known functional elements. One such region is the ribosomal frameshift site at the end of ORF1a, which has a slippery site and a pseudoknot structure [30]. Several other regions correspond to transcription regulatory sites (TRSs) [31], most of which are located in the 3′ half of the genome (Figure 6B). For instance, Figure 6C shows a region between ORF3a and E, where a sharp signature of purifying selection is directly colocated with a TRS. Interestingly, some canonical TRSs are not under strong purifying selection (motifs at the start of ORF7 and ORF8, as well as the motif towards the end of M). Both ORF7 and ORF8 are known to tolerate stop codons and do not seem to play an essential role in the viral replication in humans. There are two clearly identified regions of purifying selection that do not have an interpretation in terms of known functional elements: the region located centrally in the E gene and the region between M and ORF6.

The above fitness landscape supports our assumption that synonymous mutations can be used to estimate neutral mutation rates. There are several small regions in the genome where synonymous mutations have strongly negative fitness effects, and most of these regions co-localize with well-known functional elements. The remainder of the genome does not show evidence of purifying selection on synonymous mutations.

### Updated estimates of fitness effects of amino acid substitutions

The strategy we used to compute fitness effects (Eq. (2)) has drawbacks. The simple point estimate of the logarithm of the ratio of the observed and expected counts does not give a sense of the uncertainty associated with the estimate, and it requires a pseudocount. The uncertainty in fitness estimates stems from two sources: variation in the expected counts that is not explained by the model and sampling noise in the observed counts. The former depends on the mutation type and the context-dependent model of mutation rates reduces this variance, but the remaining variance *τ*^2^ still amounts to 3-fold variation for some mutation types (Table S1). The latter, sampling noise, is most relevant for rare mutations.

Confidence intervals of fitness effect estimates can be calculated by integrating over the uncertainty in the expected counts and the sampling noise in the observed counts (see Methods A.4). Given a mutation type, its context, and the observed counts, we have the following posterior distribution for the fitness effect of the mutation:

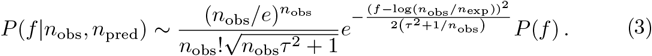

In this approximation, which is valid for moderate and large *n*_obs_, the variance of the likelihood is given by *τ*^2^ + 1*/n*_obs_. This expression highlights the contributions from the uncertainty *τ* of the underlying neutral mutation rate and of the sampling noise 1*/n*_obs_. If the prior distribution *P* (*f*) is Gaussian, then the posterior distribution is also Gaussian, and we can explicitly calculate the mean and variance of the posterior distribution (see Eq. (11)). When estimating fitness effects of amino acid mutations, which often vary considerably between neighboring sites, we use a prior that does not encourage smooth variation of fitness effects along the genome.

To assess the validity of the uncertainty estimates of the fitness, we compared pairs of mutations at the same codon that yield the same amino acid substitution. Assuming that the amino acid substitution dominates the fitness effect and that the RNA sequence of the codon has a negligible effect on fitness, the measured fitness effects of such pairs of mutations should be identical up to an error described by the posterior distribution. To check, we collected all such pairs of mutations and calculated the rescaled discrepancy as 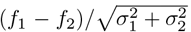, where *f*_1_ and *f*_2_ are the fitness estimates and *σ*_1_ and *σ*_2_ are the corresponding uncertainties. As expected, the discrepancy is centered near 0 and has a standard deviation of 1, indicating that the uncertainties in the fitness estimates are well-calibrated (see Fig. 7).

**Figure 7:**
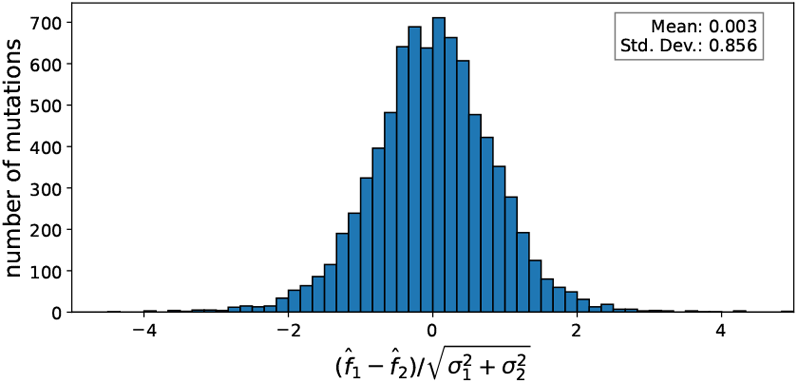
Posterior distributions of fitness estimates are well calibrated. Assuming fitness effects *f*_1_ and *f*_2_ of mutations within a codon position that lead to the same amino acid substitution are identical, their discrepancy should be centered around 0 and have a standard deviation of 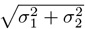. The figure shows a histogram of the rescaled discrepancy, confirming that the uncertainty in the fitness estimates captured accurately by the posterior distribution.

## Discussion

While it is well appreciated that the rate of mutation of a specific mutation type varies from position to position and depends on context, the extent of this variability in SARS-CoV-2 is remarkable. Indeed, the 5th and 95th percentiles of site-specific rates differ by ∼10-100-fold. We quantified the effects of three main features on site-specific mutation rates (genomic region, RNA secondary structure, local sequence context), and find that these effects are highly idiosyncratic between mutation types, building on Hensel’s previous work on the effects of RNA secondary structure [18]. The dependence on 3′ and 5′ sequence context reveals that some mutations are due to mechanisms that operate symmetrically on the positive and negative strand of the genome, while other mutations happen predominately on one strand. We also find evidence of both symmetric and asymmetric mutation processes, in line with previous studies [11, 32, 33]. We show that a simple model incorporating these features explains ∼ 15-60% of the fold-variation in rates depending on mutation type, and use this model to inform estimated fitness effects of mutations, identifying several small regions of the genome with strong signal of purifying selection on synonymous and non-coding mutations. At the same time, there is a substantial level of variability in synonymous mutation rates that our model is not able to explain, prompting future research into what features are missing from our model.

Although the field has identified a variety of mechanisms that lead to mutations during SARS-CoV-2 evolution, it is largely unclear which of these mechanisms are responsible for the observed patterns in site-specific mutation rates, or if some patterns arise from mechanisms that are yet unknown. At the level of genomic region, the rates of `A`→`T`, `C`→`T`, `G`→`C`, and `C`→`G` mutations all change abruptly at one of two gene boundaries along the genome sequence. To our knowledge, no previously characterized mutational mechanism is known to operate in such a way. These patterns could arise from a mechanism that acts progressively along the RNA genome before falling off, or perhaps from different regions of the genome being more or less accessible to certain mutagens. For example, the abrupt drop in the `C`→`T` mutation rate at the ribosomal slippage site could be related to translation of ORF1a and ORF1ab prior to negative-strand synthesis. ORF1a is translated more frequently than the part after the slippage site [34, 35], which could make the ORF1a RNA coding sequence more accessible to factors that cause `C`→`T` mutations. Similarly, the 3′ part of the genome is transcribed much more often than the 5′ part to produce the subgenomic mRNAs coding for abundant structural proteins like spike and the nucleoprotein [35], which could make the 3′ part more accessible to factors that cause `A`→`T`, `G`→`C`, and `C`→`G` mutations. Transcription-associated mutagenesis has been described for eukaryotic genomes [36], but it is unclear how such mechanisms would act on SARS-CoV-2.

In terms of RNA structure, four mutation types have a >2 fold difference between unpaired and paired sites. Two are associated with putative mechanisms (`C`→`T` with APOBEC3 [14, 16] and `G`→`T` with oxidative damage [13]). The other two, `C`→`G` and `G`→`C`, have not. Both of these mechanisms types likely depend on accessibility of unpaired bases and it is plausible that the resulting mutation rates are modulated by RNA secondary structure.

Local sequence context had the largest effect on mutation rate. We hypothesized that some effects would be due to specific human innate-immunity proteins that are proposed to shape SARS-CoV-2’s mutation spectrum [37] and known to preferentially target specific local sequence contexts. However, the largest effects from our analysis were not clearly attributable to known preferences of these proteins. For instance, a recent study showed that a few different APOBEC homologs were capable of inducing `C`→`T` mutations in SARS-CoV-2 RNA in cell culture, with different homologs having somewhat different motif specificities [16]. Similar to De Maio et al. [11], we find that `C`→`T` synonymous mutation counts are slightly elevated in `TCN` motifs, where `N` is any nucleotide, which aligns with motifs specificities identified in the above study [16]. However, `C`→`T` counts do not strongly depend on 3-mer sequence context (this is true for both paired and unpaired sites; Fig. S11A/B). Other studies have hypothesized that `A`→`G` and `T`→`C` mutations identified in SARS-CoV-2 genomes from patient samples and cell-culture experiments are induced by ADAR1 [38, 39]. However, the counts for these mutation types are not elevated at putative ADAR1 motifs [39, 40] (Fig. S11C). Another study found that substitutions in globally circulating SARS-CoV-2 sequences tend to be enriched at CG dinucleotides relative to other dinucleotides [37], possibly due to selection to avoid antagonism by ZAP, which targets this dinucleotide motif [41]. In our data, `C`→`T` mutation counts tend to be elevated at `NCG` motifs, but the effect is small and similar effects are not uniformly apparent for `C`→`N` and `G`→`N`. If the largest effects of local sequence context are not due to host innate immune factors, another possibility is that they are due to the viral polymerase and its associated proof-reading mechanism. Indeed, for mutation types `A`→`N` and `T`→`N`, the effects are symmetric between strands, which one might expect from a polymerase-based mechanism that copies the genome first from a positive to a negative strand and then back into a positive strand. However, the effects for `C`→`N` and `G`→`N` mutation types are not completely symmetric, and so do not lend themselves to this explanation. As the SARS-CoV-2 field continues to characterize more mutational processes, it will be interesting to see which align with patterns we characterized in this study.

The high variability in site-specific mutation rates for SARS-CoV-2 has general implications for evolutionary modeling of viruses. Classical models for detecting diversifying or purifying selection in protein-coding sequences identify codons where nonsynonymous substitutions occur more or less frequently than expected under a baseline neutral substitution process that is uniform across sites [42], with the assumption that deviation from the baseline is due to selection. However, our work on SARS-CoV-2 suggests that large deviations from the baseline can occur merely due to differences in neutral mutation rates between sites. As such, models might wrongly attribute these site-specific differences to selection. This general concern has motivated the development of models that allow for differences in neutral mutation rates between individual codon positions in a gene [43, 44, 45]. Our work broadly supports development of such models, though also highlights challenges. For instance, the above models assume that the rates of all 12 mutation types vary between codon positions in a gene in the exact same way. In some cases, this may be a reasonable assumption, and such assumptions are often necessary due to limited data. However, in the SARS-CoV-2 data, the site-to-site variability in rates has a low correlation between mutation types (in Fig. S3, most *R*^2^ values are *<*0.1).

Our study has several limitations. First, we assume that the majority of synonymous mutations are neutral, such that the number of synonymous mutation counts per site is proportional to the neutral rate of that mutation. However, synonymous mutations are not always neutral, as seen in the small genomic regions we identified where synonymous mutations have negative fitness values. And we cannot rule out the possibility that additional synonymous mutations outside these regions are under purifying selection, such that selection shapes the counts data more than we assume. Second, the ∼16 million genome sequences that we analyzed were collected using a variety of experimental and computational pipelines by laboratories around the world, which could introduce a variety of errors. Although the Bloom and Neher [24] pipeline for generating the counts data has extensive quality-control to minimize errors, and although our analysis of nonsense mutations indicated that overall error rates were low, it is still possible that errors impact the data at a subset of sites. Third, we aggregate mutation counts across the entire tree, which assumes that the mutation process is constant over time. Bloom et al.[12] found systematic but modest variation in the SARS-CoV-2’s mutation spectrum across different clades, with the largest change being a ∼2-fold decrease in the rate `G`→`T` mutations during the transition to Omicron variants (see also [13]). We verified that site-specific counts are correlated before and after the emergence of Omicron, supporting the assumption that the mutation process is largely constant over time. But the mutation process may have changed in ways that we did not capture (see Fig. S12). Lastly, the number of sites with synonymous mutation counts data was much lower for some mutation types than others (this number ranged from 261 sites for `G`→`C` and `G`→`T` to 4,430 sites for `T`→`C`), which limits our ability to resolve patterns for mutation types with less data.

In all, our study provides one of the most high-resolution analyses of a virus’s neutral mutation spectrum to date. It was already known that SARS-CoV-2’s mutation rate is highly variable between mutation types. Our finding that rates of specific mutation types are also highly variable between sites reveals substantially more heterogeneity in the virus’s neutral mutation spectrum than previously appreciated. It also raises the question of whether mutational spectra of other viruses are as variable between sites as SARS-CoV-2’s, and to what extent differences in site-specific rates depend on factors similar to the ones we identified. While it is difficult to fully answer this question at the present, as we do not have datasets of equivalent size for other viruses, our study sets the stage to make such comparisons in the future.

## A Methods

### A.1 Data and code availability

See https://github.com/matsengrp/SARS2-synonymous-mut-rate/ for data and code to curate and analyze site-specific mutation counts. Specifically, see https://github.com/matsengrp/SARS2-synonymous-mut-rate/blob/main/results/curated_mut_counts.csv for a table of curated site-specific mutation counts, which we used as input for various analyses in this paper. The README.md file in the repository’s root directory describes this file in more detail.

See https://github.com/neherlab/SARS2-mut-fitness-v2 for the updated context-aware estimates of fitness effects and associated levels of uncertainty. For each mutation, the pipeline computes a mutation’s expected counts using both the original strategy from Bloom and Neher [24] and using the log-linear model from this paper. It then separately reports estimated fitness effects for both sources of expected counts, allowing effects to be compared between these approaches. The updated context-aware fitness effects and associated levels of uncertainty are computed from the posterior distribution given by Eq. (3).

### A.2 Counting mutations on the UShER tree

As input, we used mutation counts from Bloom and Neher [24]. Specifically, we used a version of the counts that they generated by running their pipeline on an UShER tree with all sequences in GISAID as of 24 April 2024 [22, 23]. The code to run this pipeline and the resulting mutational counts are in the GitHub repository https://github.com/jbloomlab/SARS2-mut-fitness/. The pipeline generates a file called `results/expected_vs_actual_mut_counts/expected_vs_ actual_mut_counts.csv`, which reports the counts of each possible nucleotide mutation across the genome along the branches of the tree. In doing so, the pipeline divides the tree into several clades and separately reports mutation counts for each clade, only reporting counts for mutations away from a given clade’s founder sequence, as given in the file `results/clade_founder_nts/ clade_founder_nts.csv`. We then curated the counts as follows. First, we identified all sites in the genome where the nucleotide identities at that site, the site’s codon, and the site’s 5′ and 3′ nucleotides are conserved in all clade founder sequences, including the Wuhan-Hu-1 sequence (we ignore the codon requirement for noncoding sites). Next, we filtered out mutations at sites that: i) did not meet the above conservation criteria, ii) were masked in the UShER tree in any clade (see the `masked_in_usher` column), iii) were identified as error-prone (see the `exclude` column; we also filtered out the set of error-prone sites identified by De Maio et al. [27]). Next, for the remaining mutations, we summed the counts of each mutation across all clades (using the counts in the `actual_counts` column, and only summing rows where the `subset` column equals `all`, as opposed to `England` or `USA`), resulting in the site-specific mutation counts used in our analyses. To compute counts for terminal or non-terminal branches, we simply summed counts in the columns `count_terminal` or `count_non_ terminal`. To compute counts for pre-Omicron vs. Omicron clades, we summed counts for the relevant set of clades. The file https://github.com/matsengrp/SARS2-synonymous-mut-rate/blob/main/results/curated_mut_counts.csv is a table of the resulting curated site-specific mutation counts.

A large source of artificial errors in the UShER tree involve reversion mutations to the Wuhan-Hu-1 reference sequence [46]. However, we ignore reversion mutations since we only analyze mutations going away from the Wuhan-Hu-1 sequence and founder sequences (with the wildtype nucleotide identity being conserved across each).

### A.3 Randomization testing

We use randomization testing to test if the ratio of median synonymous mutation counts at unpaired to paired sites is significantly >1 (see Figure 2). For each mutation type, we randomized the unpaired/paired labels between sites and recomputed the ratio. We did this 1,000 times to generate a null distribution of randomized ratios. We computed the fraction of ratios that were greater than or equal to the observed ratio, resulting in a p-value corresponding to the probability that a value at least as extreme as the observed value is drawn from the null distribution. We multiplied the resulting p-value of each mutation type by 12 to correct for multiple-hypothesis testing via the Bonferroni method. We used a similar workflow for assessing significance in Fig. S4.

### A.4 Confidence intervals for fitness effect estimates

In Eq. (2) we provide a simple point estimate of the fitness effect of a mutation with no indication of the uncertainty associated with this estimate. The uncertainty stems from two sources: variation in the expected counts that is not captured by the model and sampling noise. To properly account for these sources of variation, we define a posterior distribution over the fitness effect *f* given the observed counts *n*_obs_ as

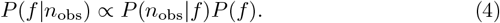

We impose a Gaussian prior 𝒩 (0, *σ*^2^) on *f* and use a likelihood that integrates over the variation in the underlying mutation rate as

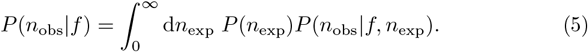

For the likelihood, we assume that

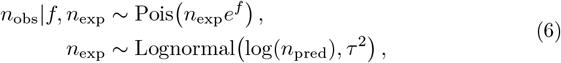

where *n*_pred_ is the mutation count predicted by the fitted model in Eq. (1) and *τ*^2^ is the remaining variance on the log-counts after the model fit for a certain mutation type. The Poisson distribution accounts for sampling variation. The likelihood is then

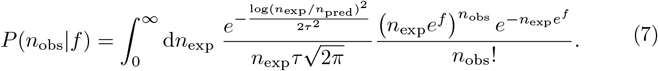

With the substitution *y* ≡ log(*n*_exp_*/n*_pred_), the expression simplifies to

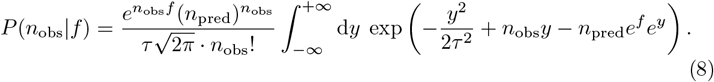

This integral cannot be solved analytically. Instead, we expand the exponent around the value of *y* where *n*_obs_*y − n*_pred_*e*^*f*^ *e*^*y*^ is maximal, *δ*_*f*_ ≡ log(*n*_obs_*/n*_pred_) *f*, to second order and eventually obtain

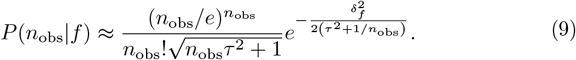

Interpreted as a function of *f*, this is a Gaussian centered around the naive estimate log(*n*_obs_*/n*_pred_) with a variance of *τ*^2^ + 1*/n*_obs_, reflecting our expectation that a more precise estimate of the underlying mutation rate reduces the uncertainty in the fitness estimate. Together with the Gaussian prior 𝒩 (0, *σ*^2^) on *f*, the above yields the approximate posterior

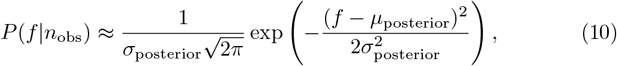

Where

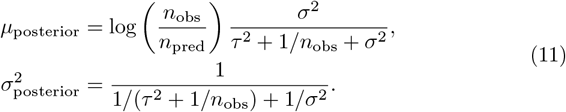

Throughout all numerical calculations we set *σ* = 2.5, which corresponds to a rather uninformative prior with a variance exceeding the variance of the naive fitness estimates across sites. While it is possible to use a more informative prior (e.g. by adjusting its mean to the empirical mean or accounting for the left-skew), the choice of prior has little effect on the posterior distribution as long as it is not too narrowly peaked. The broad zero-centered Gaussian gives rise to simple analytical expressions for the posterior mean and variance.

### A.5 Smoothed fitness estimates along the genome

Figure 6 is meant to identify positions in the genome that are under *non-coding* constraint, which is constraint that does not come from selection on protein function. Such constraint should manifest at sites of synonymous mutations and at non-coding positions outside of protein coding genes. Furthermore, we expect such constraint to affect several neighboring sites.

To detect such constraint, we sum the counts of all synonymous and non-coding mutations at a site *i* and obtain one *observed* fitness effect per site, *g*_*i*_ (undefined for sites with no synonymous/non-coding mutations). This estimate is shown for individual positions in Figure 6C. Some positions lack an estimate as there are no (non-excluded) synonymous/non-coding mutations at those sites. To smooth the trace produced by all *g*_*i*_ without introducing sliding window artifacts, we plotted in Figures 6A and 6B the *inferred* fitness effects of synonymous/non-coding mutations, {*f*_*i*_}, defined as the maxima of the following posterior

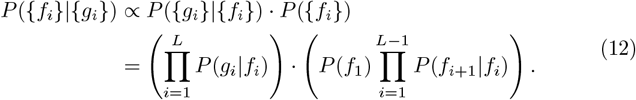

The prior encodes our belief that the degree of non-coding constraint does not vary too rapidly along the genome and the likelihood assumes that the observed fitness effect is normally distributed around its true value. Specifically, these functions are assumed to be of the form

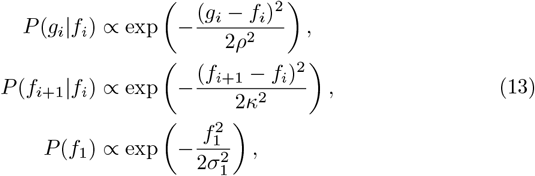

where ρ, *κ*, and *σ*_1_ are hyperparameters. Increasing the ratio *κ/ ρ* brings *f*_*i*_ closer to *g*_*i*_, while decreasing it enhances the smoothing effect. Eq. (12) can be rewritten as

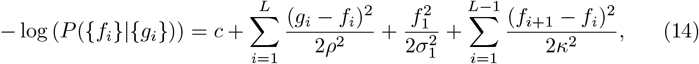

where *c* is a constant independent of {*f*_*i*_}. The condition

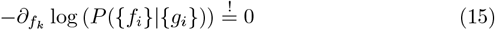

leads to the following system of linear equations:

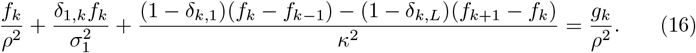

This linear system can be solved efficiently and the resulting *f*_*k*_’s constitute the smoothed fitness estimates in Figure 6. We let *ρ* tend to ∞ at all sites without at least one non-excluded synonymous/non-coding mutation, which makes Eq. (16) and *f*_*i*_ defined for all positions in the genome.

### A.6 Neural network models of mutation rate variation

We also modeled synonymous mutation counts with neural networks. These models work with similar input to the model described above and defined in Eq. (1). Genomic region and RNA secondary structure are treated as binary indicator variables. We use a more general local sequence context, which allows for reduced dimension embeddings of arbitrary fixed length motifs. Note that for a site *s* and non-negative integers *k* and 𝓁, with 𝓁 ≤ *k*, the motif centered on *s* and spanning the nucleotides at sites *s*−*k, s*−*k* +1, …, *s* +*k* is represented by the 2(*k* −𝓁) + 1 motifs of length 2𝓁 + 1 at sites *s* −*k* + 𝓁, *s* −*k* + 𝓁 + 1, …, *s* + *k* −𝓁.

For a given *k* and 𝓁, the model has an embedding of the length (2𝓁 + 1) motifs (i.e., a trainable linear layer with a parameter for each distinct motif), stacks the additional feature values for genomic region and RNA pairing state on the embedding (using binary indicator variables of each feature, as in the log-linear model from the main text), and then applies a linear layer. Fig. S10 depicts the architecture for *k* = 2 and 𝓁 = 1. This architecture is inspired by the one described in [47]. The models are trained to minimize MSE loss.

To compare the models by explained variance, we computed *R*^*2*^ values on data withheld from training. For select mutation types, complex models perform modestly better (Fig. S9). We were unable to incorporate additional layers and non-linear activation functions while improving performance, so this remains an open area.

## Acknowledgements

We thank Russell Corbett-Detig for useful discussions and Angie Hinrichs for her help with UShER trees, useful discussions, and her heroic effort to maintain and grow pandemic-scale trees of all available SARS-COV-2 data. We gratefully acknowledge all data contributors, i.e., the authors and their originating laboratories responsible for obtaining the specimens, and their submitting laboratories for generating the genetic sequence and metadata and sharing via the GISAID Initiative, on which this research is based.

