## Supplementary Information for "The mutation rate of SARS-CoV-2 is highly variable between sites and is influenced by sequence context, genomic region, and RNA structure"

January 7, 2025

### Figures and Tables

| mutation type | number of sites | remaining variance $\tau^2$ |
|---|---:|---:|
| A→C | 1,479 | 0.519 |
| A→G | 2,426 | 0.227 |
| A→T | 1,291 | 0.669 |
| C→A | 649 | 0.488 |
| C→G | 532 | 0.886 |
| C→T | 1,539 | 0.476 |
| G→A | 862 | 0.350 |
| G→C | 261 | 1.167 |
| G→T | 261 | 0.971 |
| T→A | 2,280 | 0.783 |
| T→C | 4,430 | 0.276 |
| T→G | 2,032 | 0.618 |

**Table S1: Characteristics and summary statistics of different mutation types.** Column 1 lists the number of sites that met the criteria described in Figure 1A and are used in the estimation of the model. Column 2 shows the remaining variance in logarithmic count of mutations at these sites that is not explained by the model. tab:n sites

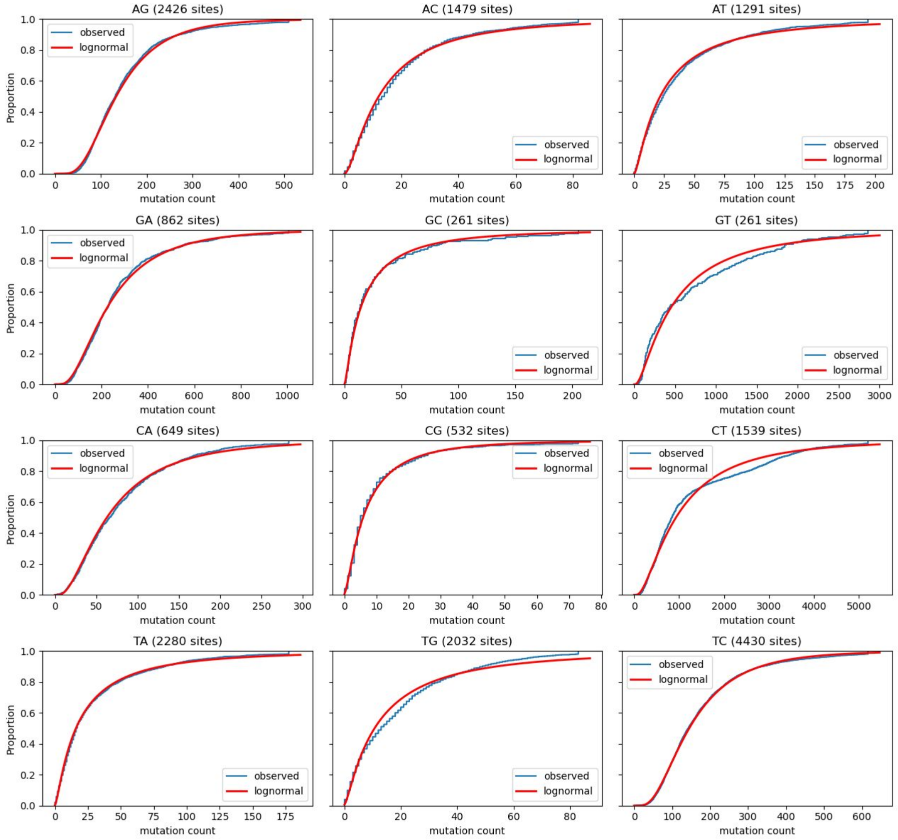


**Figure S1: For each mutation type, the synonymous mutation counts per site closely follow a log-normal distribution.** In each plot, the blue curve shows the cumulative distribution of the observed synonymous mutation counts per site for a given mutation type, while the red curve shows the cumulative distribution of a log-normal distribution with the same mean and variance as the corresponding observed distribution. In each case, the blue curve closely tracks with the red curve, with only a few deviations, showing that the observed counts closely follow a log-normal distribution.

fig:log˙normal

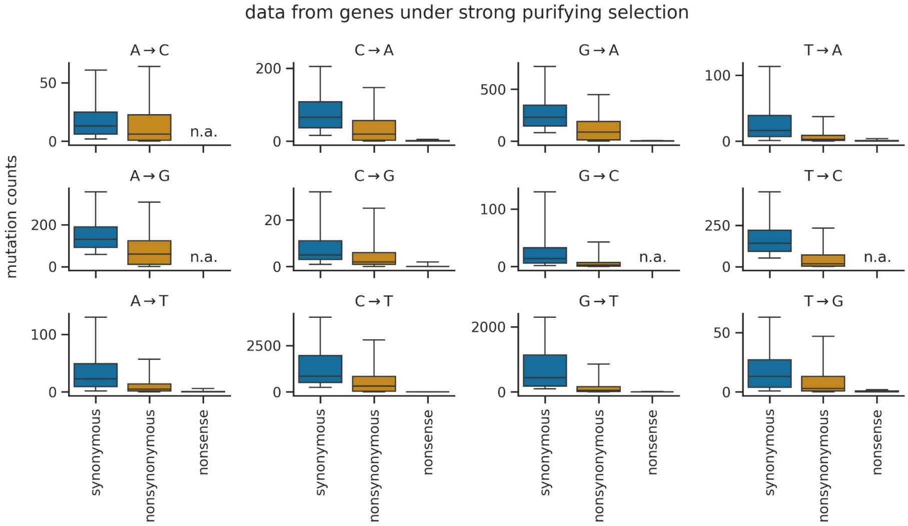


**Figure S2: Distributions of site-specific mutation counts for synonymous, nonsynonymous, and nonsense mutations for each mutation type.** To examine the possibility that artificial errors could impact the counts, we computed site-specific mutation counts for nonsynonymous and nonsense mutations using the same basic approach that we used for synonymous mutations. For each mutation type, box plots show site-specific mutation counts for synonymous, nonsynonymous, or nonsense mutations (whiskers show the 5th and 95th percentiles). Plots only show data for mutations in genes under strong purifying selection according to [1] (ORF1a, ORF1b, S, E, M, N), which make up most of the genome. Given the structure of the genetic code, some mutation types never lead to a stop codon in any context, and the plots for these mutation types have "n.a." instead of counts data for nonsense mutations. If the observed counts of *synonymous* mutations are inflated by artificial errors, then we would expect counts of nonsynonymous and nonsense mutations to be inflated by the same amount, as artificial errors are not subject to natural selection. On the other hand, if the observed mutation counts actually correspond to mutations that arose in globally circulating viruses and have been subjected to natural selection, we would expect the counts for nonsynonymous and nonsense mutations to be lower than counts for synonymous mutations, as nonsynonymous mutations are often deleterious and nonsense mutations are almost always highly deleterious. Overall, the distributions are consistent with the expected action of natural selection. Notably, nearly all nonsense mutations have counts near zero, indicating that artificial errors probably do not substantially inflate synonymous mutation counts in most cases. fig:mut˙counts˙supp

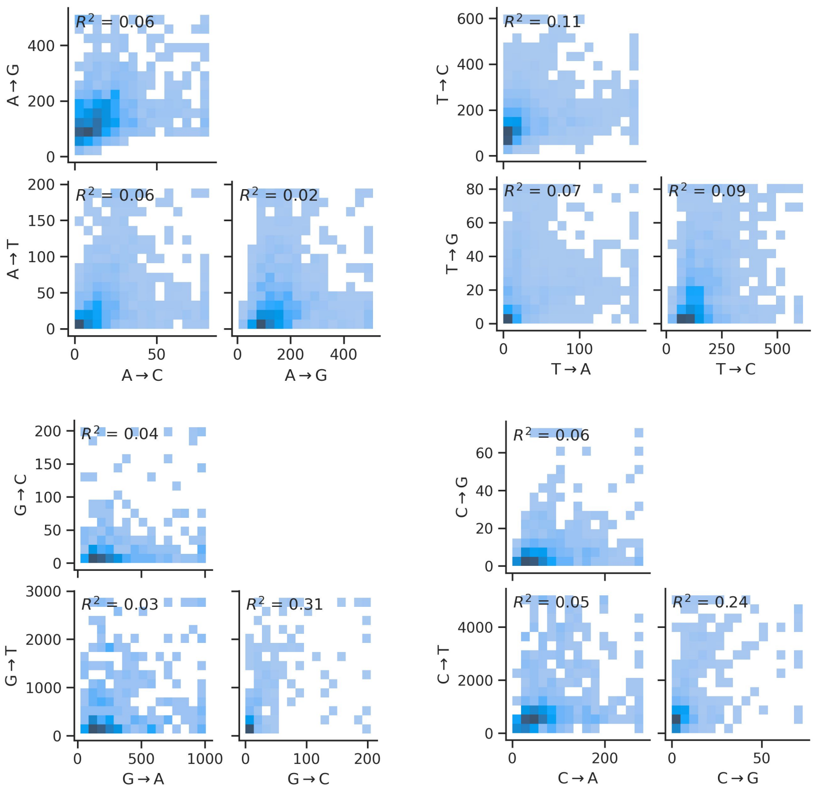


**Figure S3: Correlation of synonymous mutation counts between different mutation types at the same site.** For each mutation type, we apply a ceiling to the counts distribution at its 98th percentile. For most mutation types, the correlation is low, with $R^2 < 0.1$. Though, for a few pairs of mutation types (G→T and G→C or C→T and C→G), the correlation is substantially higher, indicating that counts of these pairs are more strongly influenced by common factors. One factor might be RNA structure, as these are the four mutation types where RNA structure has the largest effect on counts (Figure 2E).

fig:mut˙type˙corr

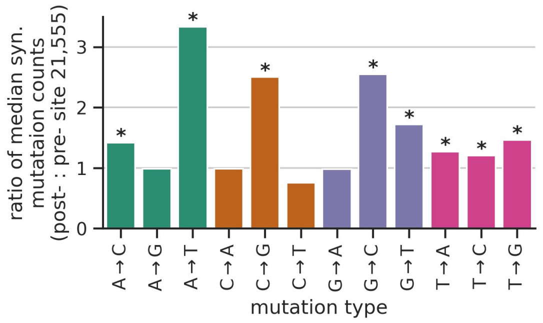


**Figure S4: Position dependence of A→T mutation counts is also observed for C→G and G→C.** The plot shows the ratio of median synonymous mutation counts between sites before vs. after site 21,555 in the genome. Asterisks indicate mutation types where the ratio of medians is significantly >1.0, as determined by randomization testing (see *Methods*), using a threshold of $p<0.05$ after Bonferroni correction for multiple-hypothesis testing. The largest effect sizes are for A→T, C→G and G→C. Some of the other mutation types show significant effects, but the effect sizes are not as large.

fig:genomic˙region˙pattern

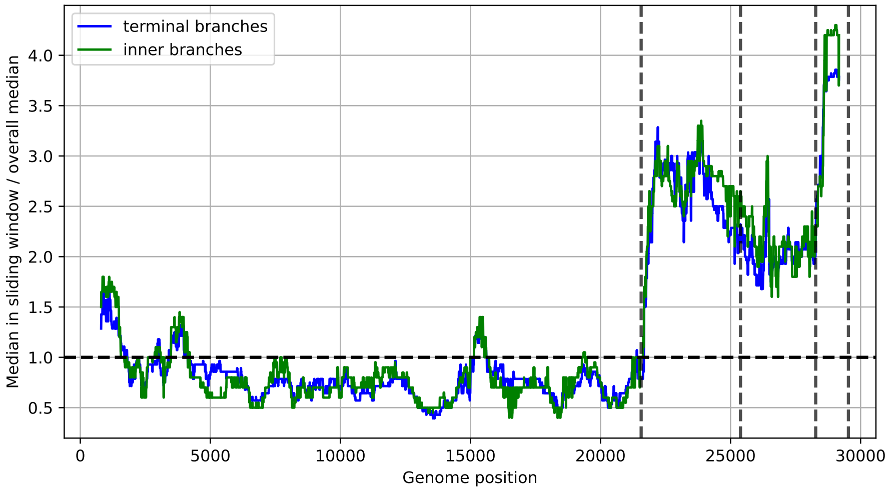


**Figure S5: Position dependence of A→T mutation counts is observed for terminal and non-terminal mutations.** The sharp increase of A→T mutation counts at the boundary between ORF1ab and S is not restricted to terminal mutations. Going from left to right, the first and second vertical lines mark the beginning and end of the S coding sequence, while the third and fourth vertical lines mark the beginning and end of the N coding sequence.

fig:AT˙terminal˙nonterminal

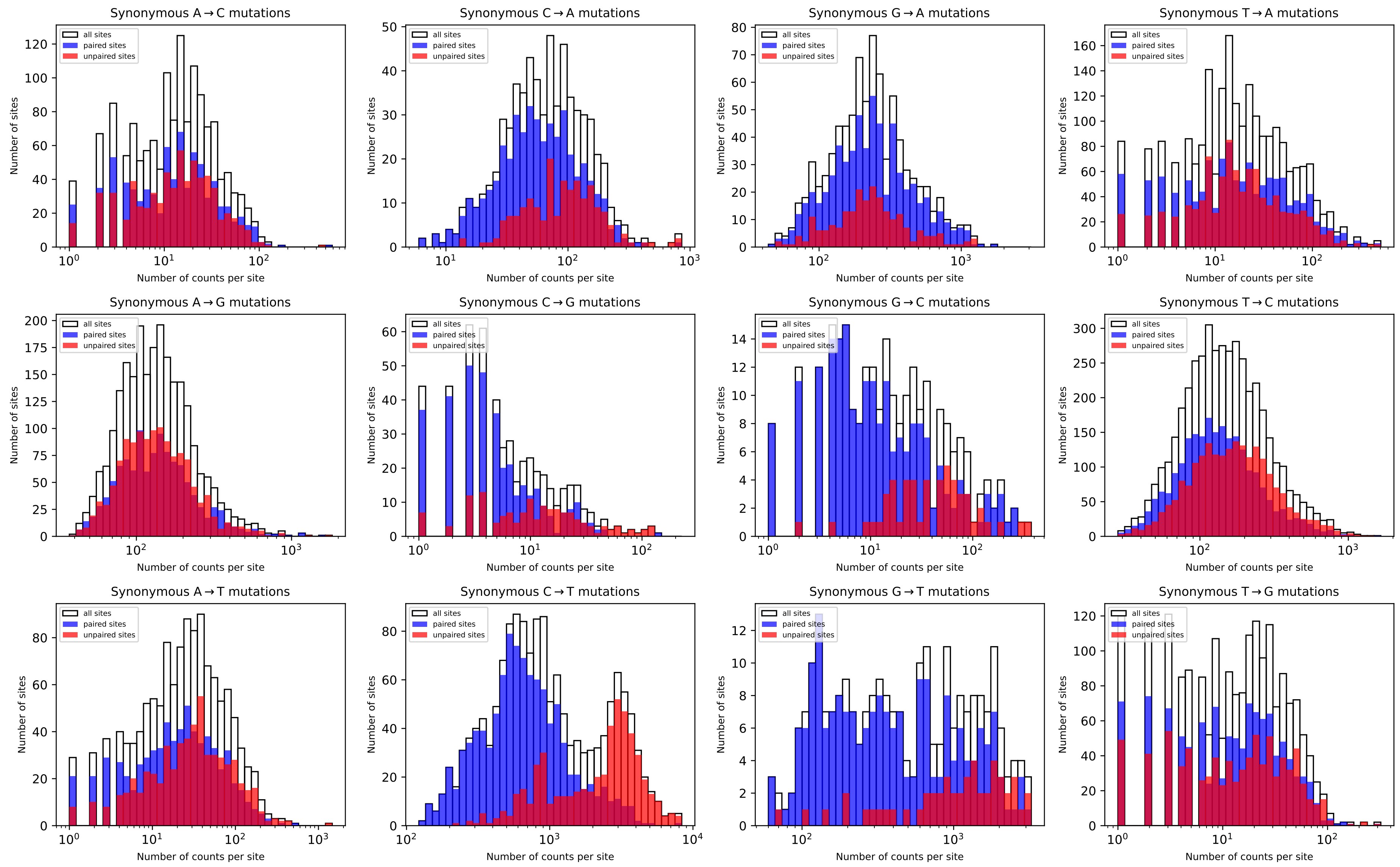


**Figure S6: Distributions of mutation counts at sites that are base-paired or unpaired in the predicted secondary structure of the SARS-CoV-2 genome.** Each panel shows the distribution of the mutation counts split by pairing state. Note that for some mutations, C→T in particular, the distribution of counts at unpaired sites is shifted to higher values compared to that unpaired sites.

fig:paired˙unpaired

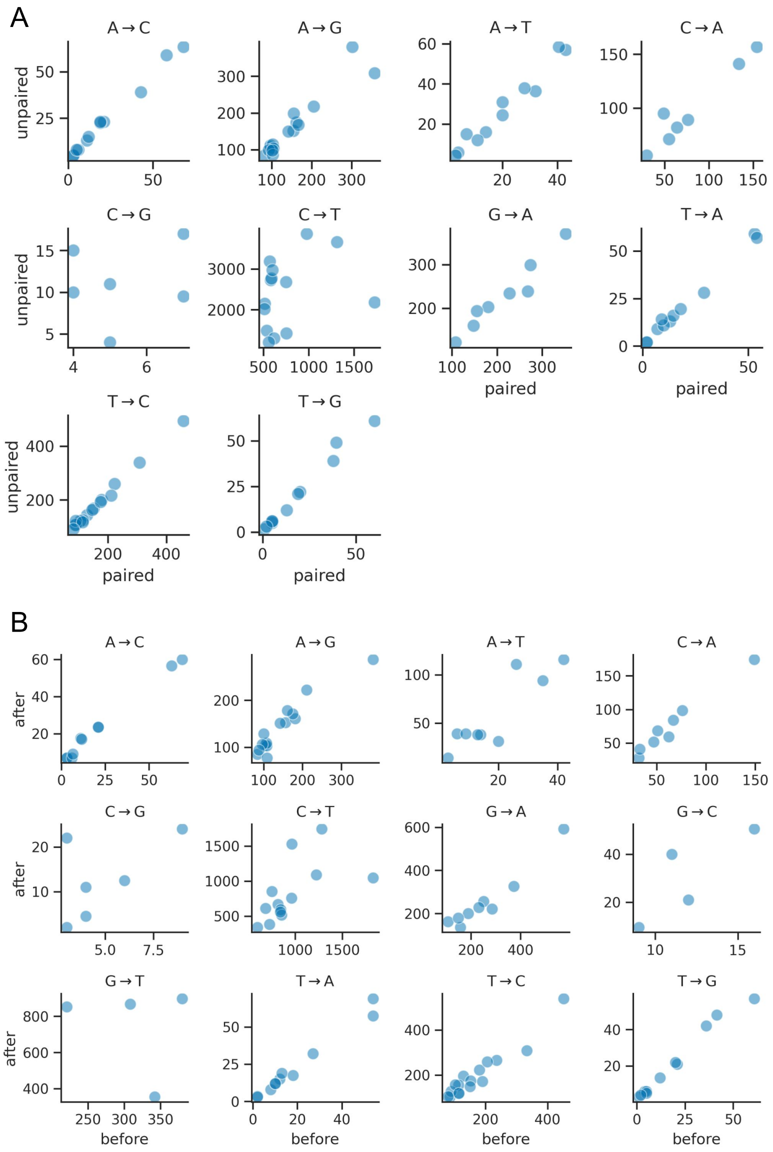


**Figure S7:** Correlation of 3-mer motif median synonymous mutation counts between subsets of sites in the genome for a given mutation type. **(A)** Comparing sites that are paired vs. unpaired in the predicted secondary structure. **(B)** Comparing sites before vs. after nucleotide 21,555 in the SARS-CoV-2 genome, which marks the end of ORF1b, the point at which synonymous mutation counts for A→T increase dramatically. These plots only show data for motifs with at least 10 sites in each subset for a given mutation type (a few mutation types do not have any motifs meeting this criterion).

fig:local˙context˙differences

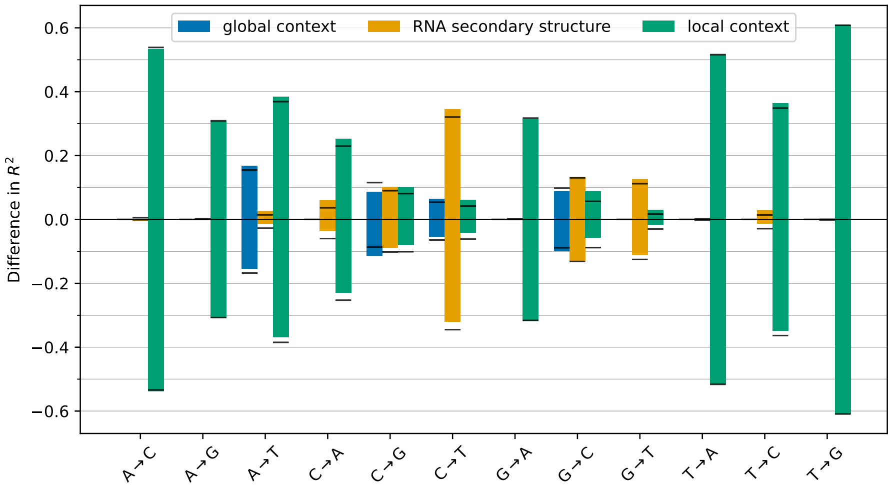


**Figure S8: Individual contributions of different features are largely independent.** The bars show the increments and decrements of the explained variance when features are used in isolation or removed from the full model, respectively. The horizontal black lines show the value of each bar mirrored across the x-axis. In almost all cases, the decrement upon removal matches their individual contribution, showing that these features are non-redundant.

fig:separately˙R2

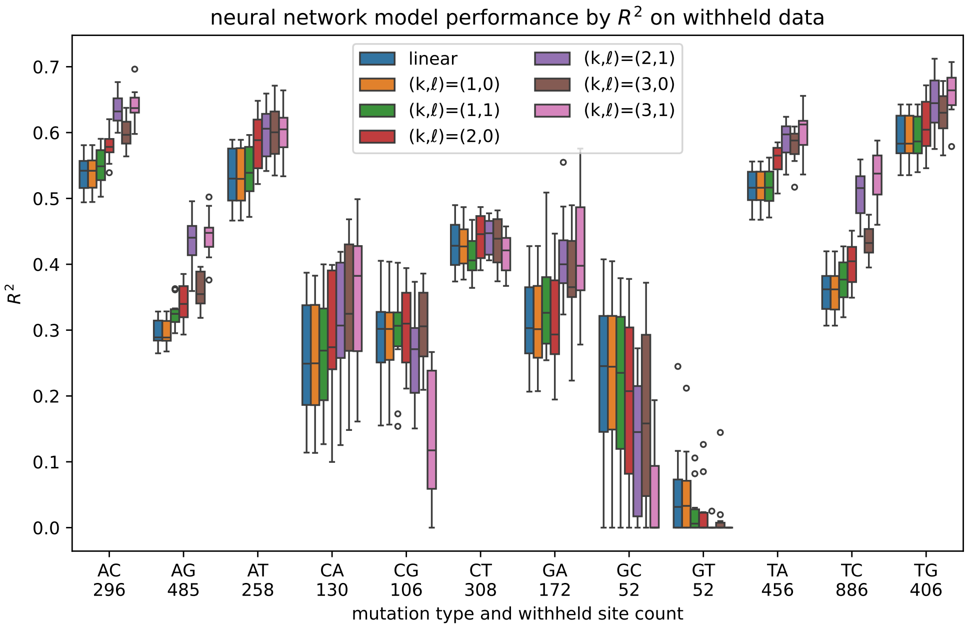


**Figure S9: Performance of neural-network models and the linear model from the main text in their ability to predict site-specific synonymous mutation counts.** We devised a neural-network architecture for predicting counts based on a site's genomic region, RNA structure, and local context (see *Methods*). We examined a series of models based on this architecture that integrate information over larger and larger windows of local sequence context (Fig. S10). The legend refers to these models using the convention described in the *Methods* and Fig. S10 (models with $k$ of 1, 2 or 3 integrate information across sequence windows of length 3, 5, or 7, respectively). The "linear" model in the legend refers to the linear model from the main text. For each mutation type, we trained each model using a random 80% of sites and tested the model using the withheld 20% of sites, computing the $R^2$ between the predicted and observed counts from the withheld data. We did so using 10 randomly generated train-test splits, and each box plot shows the distribution of the resulting 10 $R^2$ values. For mutation types with more data (see the x-axis for the number of withheld sites per mutation type), the more complicated neural-network models (larger $k$ and $\ell$ values) tended perform better than the linear model (higher $R^2$). However, for mutation types with less data, their performance was worse, suggesting more overfitting. Even when the neural networks performed better, the linear model still captured most of the variance captured by the neural networks, prompting us to use the linear model for estimating fitness effects. fig:nn`model`r2

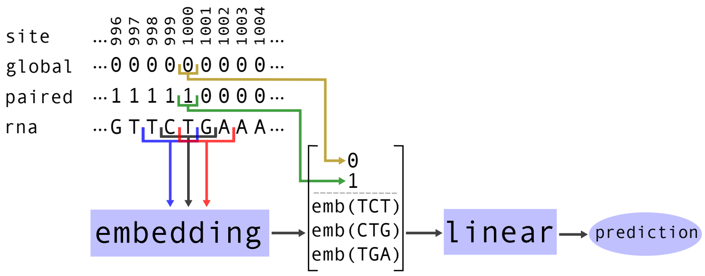


**Figure S10: Summary of neural-network architecture.** This figure shows an architecture where $k = 2$, $\ell = 1$ (see *Methods*), and how this architecture integrates information about local sequence context, RNA pairing state, and genomic (global) region, for site 1000. In this example, the length 5 motif centered at site 1000 is represented by the length 3 motifs centered at sites 999, 1000, and 1001. fig:nn`model`architecture

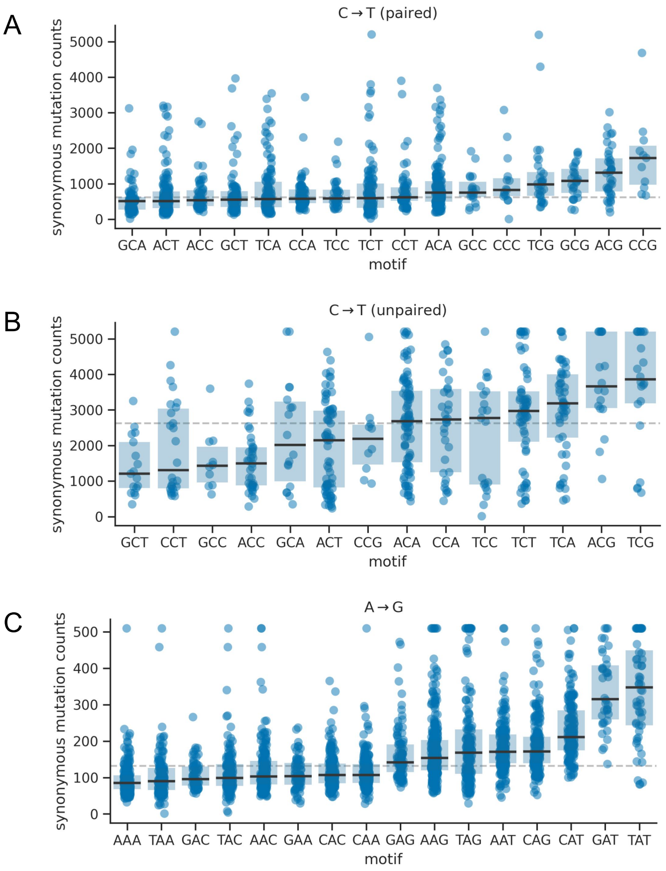


**Figure S11: Effect of local sequence context on synonymous rates for additional mutation types.** The panels in this figure are similar to the ones from Figure 3A/C. **(A)** and **(B)** separately show data for C→T mutations at sites that are paired or unpaired in the predicted secondary structure of the SARS-CoV-2 genome. For both sets of sites, there are only modest differences in the median mutation counts between different 3-mer motifs. **(B)** shows data for A→G mutations. A previous study identified putative ADAR1 motifs as being depleted in G nucleotides one base upstream of the mutated site and enriched in G nucleotides one base downstream of the site [2]. Another study identified the motifs as also being enriched in A nucleotides one base upstream and enriched in A nucleotides one base downstream [3]. These putative motifs do not clearly correspond to the ones with the highest counts. fig:local context ct ag

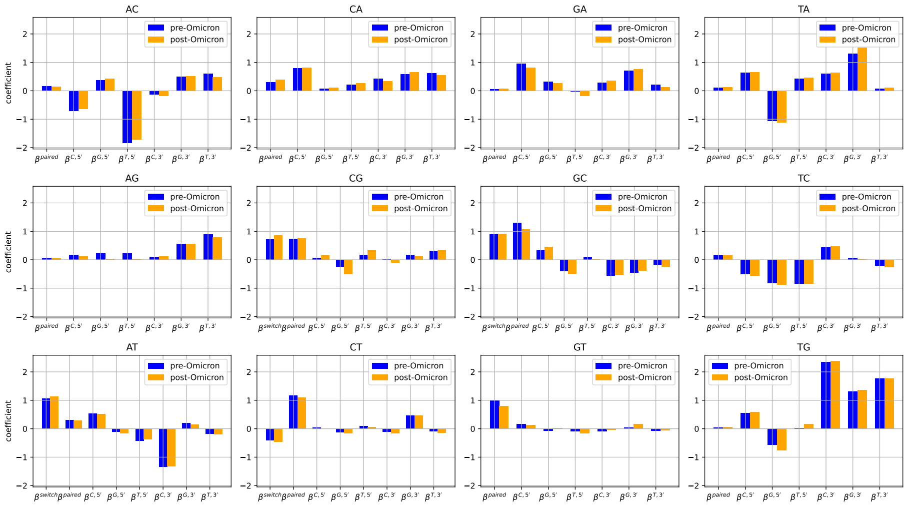

**Figure S12: Comparison of model parameters inferred from data before and after the transition to Omicron variants.** The effect of sequence context, RNA secondary structure, and large scale genomic regions is very similar for Omicron and pre-omicron variants.

fig:pre˙vs˙omicron
